# Neural processing of iterated prisoner’s dilemma outcomes indicates next-round choice and speed to reciprocate cooperation

**DOI:** 10.1101/784033

**Authors:** Francisco Cervantes Constantino, Santiago Garat, Eliana Nicolaisen-Sobesky, Valentina Paz, Eduardo Martínez-Montes, Dominique Kessel, Álvaro Cabana, Victoria B Gradin

## Abstract

Electing whether to cooperate with someone else is well typified in the iterated prisoner’s dilemma (iPD) game, although the neural processes that unfold after its distinct outcomes have been only partly described. Recent theoretical models emphasize the ubiquity of intuitive cooperation, raising questions on the neural timelines involved. We studied the outcome stage of an iPD with electroencephalography (EEG) methods. Results showed that neural signals that are modulated by the iPD outcomes can also be indicative of future choice, in an outcome-dependent manner: (i) after zero-gain ‘sucker’s payoffs’ (unreciprocated cooperation), a participant’s decision thereafter may be indicated by changes to the feedback-related negativity (FRN); (ii) after one-sided non-cooperation (participant gain), by the P3; (iii) after mutual cooperation, by late frontal delta-band modulations. Critically, faster choices to reciprocate cooperation were predicted, on a single-trial basis, by P3 and frontal delta modulations at the immediately preceding trial. Delta band signaling is considered in relation to homeostatic regulation processing in the literature. The findings relate feedback to decisional processes in the iPD, providing a first neural account of the brief timelines implied in heuristic modes of cooperation.

## Introduction

Organized social life is based on balancing self and group benefit-seeking, a hallmark of cooperation^1^. Social scientists often model sustained cooperative interactions based on the iterated prisoner’s dilemma (iPD)^2,3^. In this iconic game, two agents make simultaneous and independent decisions on whether to cooperate or defect with each other, and each agent receives a payoff depending jointly on both decisions. One-sided outcomes lead to imbalanced payoffs that may trigger reports of anger or betrayal in the cooperator and elation (but also guilt) in the defector; balanced mutual cooperation usually induces positive outcomes such as bonding or trust^4,5^. The iPD outcomes imply varied social contexts that allow to recreate relevant scenarios from which to investigate the neural dynamics of cooperation. Through iPD outcomes’ analysis it is also possible to explore how different social contexts relate to future decisions on cooperation.

Cooperative cognition and its neural bases have been addressed in relation to feedback processing at the iPD, typically via functional magnetic resonance imaging^4,6–14^ and in some cases including populations of clinical interest^5,15–21^. High temporal resolution electroencephalography (EEG) methods have also been applied^22–24^, with two previous studies directly addressing the processing of the game outcomes^25,26^. These two studies identified two relevant event related potential (ERP) components, the feedback related negativity (FRN) and the P3 (or P3b), both of which are often featured in the feedback processing literature^27^. The FRN is a deflection elicited shortly after feedback (200 to 350 ms) and represents the earliest index in reward outcome evaluation; it usually shows a more negative amplitude following negative versus positive outcomes^27–33^. When social feedback is involved, the FRN may show sensitivity to social comparisons^34^, fairness^35–38^ and cooperation^25,26^, and may comprise the affective impact of an event^28,39^. In *value-based* decision-making studies, the P3 (or P3b) is a mid-latency (300 to 600 ms) centroparietal deflection showing sensitivity to reward magnitude, valence and probability or risk^27,30,40–48^. In social studies it has similarly shown modulation by social comparisons^34,49^, fairness^37,49,50^ and cooperation^26^.

ERP analyses are popular tools for inferring neural dynamics and timing of economic decision-making, and so are spectrotemporal methods^25–27, 51^. However, to our knowledge, no study has investigated yet the late (e.g. > 600 ms) ERP signals elicited by the iPD outcomes, neither the associated frequency band activity. Feedback ERPs often include the late positive potential (LPP, cf.^52^) which is involved in emotional processing^27,53–56^. Spectrotemporal activity involved in feedback processing (e.g.^57^) has not been addressed in the game, except for derivate measures in hyperscanning protocols^22–24^. The relatively late window (> 500 ms) represents an intermediate phase between early neural updating by the outcome and next round decision-making behavior. Therefore, an investigation of feedback processing related to decision making may need to address these later processes, which also potentially mediate integration of the slow dynamics sometimes involved in decision-making^27^.

The current study had two main objectives. First, we set to comprehensively characterize EEG signals from the feedback processing stage of the iPD, including an analysis of FRN and P3 but also of (previously unexplored) late ERP signals and corresponding spectrotemporal activity. As a second objective, we investigated how the social context imposed by each outcome type relates to next decision. At this point, deciding whether to cooperate relies on potentially different assessments about the co-player, e.g. whether they are interested in a sustained, retaliatory, or deceptive interaction. This leads to situations where, even after experiencing the same outcome (e.g. CD’s on separate occasions), a player may choose to cooperate or not thereafter. For this we investigated, for each outcome type, whether the same neural signals demonstrating sensitivity to outcomes were also predictive of a participant’s subsequent choice. Of particular interest was the potential timing relationship between relevant neural activity after outcome presentation and effective choice at the next round. Time is a central variable in the ‘social heuristics hypothesis’ (SHH), a dual-process theory^58,59^ application to cooperation which distinguishes fast, intuitive decisions from ‘second thoughts’ of a more self-interested nature^60,61^. Recent studies suggest that systematically ‘defaulting’ on cooperative choices may be linked to fast, heuristic decision-making^62,63^. Hence, we employed the present feedback signal and timing analyses as a means to address candidate neural correlates of automatic decision-making when facing the prospect of repeated cooperative interactions.

## Materials and methods

### Subjects

Thirty-one volunteers (16 female; mean age 22.3 ± 2.9 SD) with no history of neurological or psychiatric disorders participated in the present study. All reported normal or corrected-to-normal visual acuity, and provided written informed consent for their participation. All experiments were performed in accordance with WMA Declaration of Helsinki guidelines. Approval of the experimental procedures was obtained by the Faculty of Psychology at Universidad de la República.

### Experimental setup

Presentation and response time logging were performed with PsychoPy^64^ software, delivered over a CRT monitor (E. Systems, Inc., CA) with 40 cm size, 83 dpi resolution, and 60 Hz refresh rate. EEG recordings were performed using a BioSemi ActiveTwo 64-channel system (BioSemi, The Netherlands) with 10/20 layout, at 2048 Hz digitization rate with CMS/DRL (ground) and two mastoid reference electrodes. A 5^th^ order cascaded integrator-comb low-pass filter with −3 dB point at 410 Hz was applied online, after which signals were decimated to 256 Hz. Online high-pass response was fully DC coupled. Full experimental sessions lasted ~2.5 h.

#### Iterated prisoner’s dilemma task

Each participant was introduced to a same-sex person, presented as their co-player for the study but in fact a confederate, i.e. an associated researcher with no prior knowledge of the participant. The confederate and the participant were then escorted to different rooms. The participant was instructed that, on each trial, both players would have to make a simultaneous and independent decision regarding whether to cooperate or not cooperate with each other. On each round, and depending on their decisions, they would both receive points. Mutual cooperation (‘CC’) [non-cooperation (‘DD’)] earned both 2 [1] points respectively, while unbalanced outcomes (‘CD’ and ‘DC’) earned 0 points for the player that cooperated and 3 for the player that did not. Participants were told that points accumulated over time, and were instructed to maximize earnings. They did a practice test and waited for the experimenter to supposedly provide similar instructions to the confederate.

During EEG recordings, participants played the iPD over a 14.4° by 6.2° visual interface. In each round, the payoff matrix was displayed (Figure 1A) and participants were required to decide whether to ‘Cooperate’ (‘C’) or ‘Not cooperate’ (‘D’) by pressing one of two buttons (left/right). After a fixation, the “joint” final outcome was highlighted as feedback, e.g. a ‘CD’ outcome means the participant cooperated but the co-player did not (i.e. a ‘sucker’s payoff’). Decision-making by the confederate was simulated by a probabilistic ‘tit-for-tat’ style algorithm. In the first round, the algorithm cooperates and in subsequent rounds it reciprocates the player’s choice with 80% probability; after three consecutive rounds of identical, mutually-balanced outcomes, it switches to the alternative option (i.e. after three CC outcomes, the algorithm switches to D). Three pauses were offered every 50 rounds, completing the session with 200 rounds. Afterwards, participants rated their emotional response (happiness, anger, sadness, betrayal, and guilt) to each of the four outcomes on 8-level Likert scales in order to address recalled emotional impact of the game setup. Participants were debriefed and one subject reported disbelief of the cover story being excluded from the analyses. Thus, the final sample for analyses included 30 subjects. Participants received a cinema ticket in appreciation for their time. In one subject’s EEG recording, data transfer was incomplete for the first half of the session and the remainder was included in the analyses.

**Figure 1.**
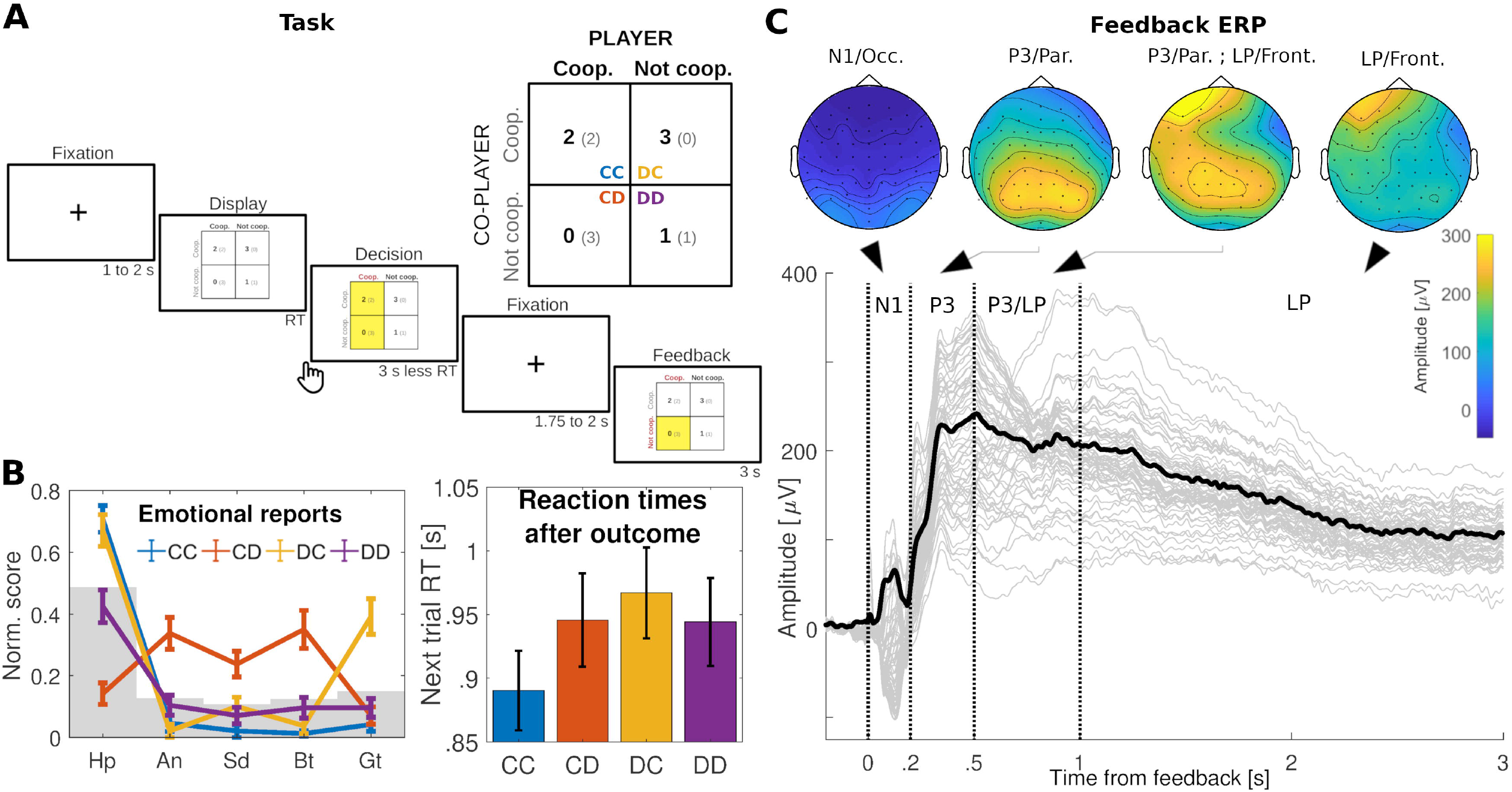
iPD presentation and associated emotional, behavioral and feedback signals. (A) Visual display of the task. The game matrix (inset) assigns earnings to both players as a joint function of their decisions to cooperate with each other. Resulting combinations apportion different amounts to player vs. co-player earnings (indicated in large vs small fonts respectively). CC: mutual cooperation; DD: mutual non-cooperation; CD: one-sided cooperation/’sucker’s payoff’; DC: one-sided non-cooperation). At each trial, the matrix is displayed after a fixation period, prompting for the participant’s decision. Given his/her choice, the two possible outcomes are briefly highlighted. After a second wait period, the final outcome is indicated. (B) Left: across participants (N=30), each outcome type leads to a distinct emotional profile (Hp: happiness; An: anger; Sd: sadness; Bt: betrayal; Gt: guilt). Grey level indicates outcome report averages by emotion. Right: Grand average of next-round reaction times at the iPD are outcome-dependent. Responses after mutual cooperation are on average faster than after one-sided non-cooperation. Error bars indicate ±1 SEM. See Supplemental Data for emotional and behavioral analyses. (C) In the iPD, feedback signals begin with occipital N1 activity peaking around 100 ms, followed by a fast-rising parietal P3 component which vanishes until about 1 s. Transition into a frontal, left-lateralized late potential (LP) begins at about 500 ms, and is sustained until the end of the trial. Topographies indicate time averages over the selected windows. Black line indicates root-mean-square over channel data (grey).

### Data analysis

#### EEG preprocessing

EEG data were referenced offline to the mastoid average and DC offset was removed. A fourth-order 0.1 – 30 Hz Butterworth filter was applied in the forward and reverse direction. Trials were epoched - 0.25 to 3 s re to outcome presentation. To remove signal artifact, single-trial and -channel data were rejected by a variance-based criterion^65^ across all channels and epochs (confidence coefficient = 4), resulting in removal of 1.75% of channel-epoch timeseries across the experiment (subject range 0.1% - 13.9%).

Feedback signals’ spatial topographies were obtained after averaging epochs across all outcomes and participants. In this grand average (Figure 1C), a sequence of N1, P3 and a frontal left-lateralized late potential (‘LP’) was identified, approximately corresponding to interval windows 0 – 0.2 s, 0.2 – 0.5 s, and 1 – 2.5 s, respectively. To prevent experimenter bias in sensor selection^27,66^, a joint decorrelation spatial filtering procedure^67,68^ was applied. The resulting set of three (non-orthogonal) linear combinations maps the original 64-channel dataset into 3 feedback components (Supplementary Figure 1A,B) that, separately, optimize detection of evoked activity within their respective time intervals. To address the feedback-related negativity (FRN) component, the spatial filtering procedure was applied to the P3 dataset (the 0.2 – 0.5 s interval after outcome onset), which contains the classic FRN window^34,40,69^. Due to relative amplitude difference with respect to the P3 in the ERP grand average, spatial filtering was applied to extract reproducible activity representing the FRN in terms of a difference between two conditions in repeated trials^68^. We estimated the subspace of the P3 dataset which maximized the difference between specific mutual cooperation (CC) and unreciprocated cooperation (CD) outcomes. As seen in emotional data, both conditions represented the most and least emotionally rewarding outcomes in the iPD from the player perspective, respectively. CC and CD trial waveforms were projected onto the resulting spatial filter.

#### Feedback signal statistical analysis

Data from participants attaining ≥ 10 trials per outcome type were included in the analysis (*n*=26). N1, FRN, P3, and LP feedback signals were window-averaged according to the corresponding epoch in which the underlying evoked component was observed to be present, as indicated above. General feedback components (i.e. N1, P3, and LP) were analyzed by the separate decisions that the player and the co-player made leading to the presented outcome. As with behavioral data, averages were submitted to a 2-way repeated measures ANOVA, with Player and Co-player as within-subject factors. FRN differences between CC and CD outcomes were analyzed by a Student’s *t*-test.

#### Spectrotemporal analysis

Evoked power from iPD feedback conditions and components was obtained from spectrotemporal correlograms over a log-spaced frequency step range of 1–25 □ Hz (200 spectral bins with power-5 spacing), via the continuous Morlet wavelet transform, log-transformed to dB-scale and referenced to average baseline values. This corresponds to the event-related spectral perturbation^70,71^ estimate of the ERP. For each feedback signal, corresponding time-frequency maps were analyzed within windows as indicated above. Spectrotemporal data were transformed to spectrotemporal *t*-map difference contrasts between outcome condition pairs, and resulting *t*-clusters above an a priori *t=2* threshold were submitted to nonparametric statistical testing^72,73^ with *N*=1024 resamplings. To avoid systematic baseline differences by player’s foreknowledge of his/her executed choice, contrasts only involved conditions with a common participant action (‘CC’ versus ‘CD’, ‘DC’ versus ‘DD’). After significant clusters were identified, spectrotemporal measures were estimated by double integration of spectrotemporal power across the time and frequency boundaries given by the cluster. Spectral domain integration was performed in linear scale. All statistical analyses were performed with MATLAB^®^ (The Mathworks, Natick, MA) except where indicated.

### Next-decision analyses

#### Feedback timeseries and spectrotemporal cluster data

In this stage of the study we focused on feedback timeseries and/or spectrotemporal cluster results that reflected modulations by outcome processing. FRN, P3, LP and LP-delta feedback signals (see Results) were tested for the hypothesis that they may relate to the participant’s decision at the next round. For each signal (e.g. FRN) and outcome type (e.g. CD), data were partitioned by whether at the next round the player would cooperate or not (e.g. ‘C|CD’ versus ‘D|CD’, respectively) in order to reveal strategic social decision-making differences stemming as early as during outcome processing. As before, data from participants attaining ≥ 10 trials per condition (*n*=16 for C,D|CC; *n*=18 for C,D|CD; *n*=21 for C,D|DC) were analyzed. Data were tested for normality by Shapiro-Wilk tests implemented with SPSS Statistics 24 (IBM Corporation, Armonk NY), and submitted to Student’s *t*, or alternatively to Wilcoxon rank-sum testing as applicable. After these analyses, to discard the possibility that any observed effects may be due to signal-to-noise differences from imbalanced sample sizes (e.g. more CC trials leading to cooperation than not), additional non-parametric randomization tests were performed^74^. For each contrast, e.g. ‘C|CC’ versus ‘D|CC’ trials, subject data were randomly shuffled with resampled trial groups maintaining the original imbalance sizes (i.e. resampled C|CC trials with a same size as the empirical sample). In each experiment resampling, shuffling was done independently per subject, and the testing randomization distribution consisted of 2^15^ experiment resamplings. Significance of empirical effect sizes was assessed by one-sided testing of the hypothesis of no difference between next-choice conditions.

#### EEG and reaction time correlations

Feedback signals found to be sensitive to next-choice conditions (P3 and LP-cluster, see Results) were examined at the single-trial level for associations to forthcoming reaction times. To test the strength of the linear association between feedback signal and next reaction time, single trial data (*n*=26 subjects) were submitted to repeated measures Pearson correlation analysis^75^ accounting for within-subject non-independence of trial-based measures. Repeated measures correlation tests were implemented with RStudio, version 1.2.1335 (RStudio, Inc., Boston, MA).

## Results

### EEG signals of iPD outcome processing

The iPD outcomes elicit a sequence of N1, P3 and a left-lateralized frontal late (‘LP’) potential (Figure 1C). A 2 x 2 factorial repeated measures ANOVA of the N1 window average amplitude with Player and Co-player choices as independent variables, showed no significant main effect of Player or Co-player choice nor an interaction (Figure 2A,B). For P3, a significant main effect of Player choice (*F*(1,25)=9.74; *p*=0.005; η_p_^2^=0.28) and a significant main effect of Co-player choice (*F*(1,25)=11.94; *p*=0.002; η_p_^2^=0.32) were found, but no significant interaction. Choices to cooperate led to greater P3 window average amplitudes (Figure 2A,B). LP revealed a significant main effect of Co-player choice (*F*(1,25)=4.68; *p*=0.040; η_p_ =0.16), but not of Player choice and no significant interaction. As before, cooperation led to greater LP window averages (Figure 2B). In addition, the outcome-specific FRN signal, which showed a centroparietal scalp topography (Figure 2C), revealed a significant difference in its window average for CC (2.0 ± 2.1 ad.u.; mean ± SD) versus CD (−0.2 ± 2.1 ad.u.) outcomes (*t*(25)=5.55; *p*<0.001; Cohen’s *d*=1.09). Spectrotemporal evoked power of the FRN, P3 and LP was separately addressed, to test for modulations of these feedback signals not directly reflected on their waveforms. Of these signals, only the LP was significantly modulated by outcome type in evoked spectrotemporal analyses. Slow-wave activity clusters within the delta band (termed ‘LP-delta’), showed a significant difference for the CC versus CD mean spectrotemporal power, in the 0.6–2.2 s (low-delta, *p*=0.002), and 1.6–2.5 s (high-delta, *p*=0.034), respectively. In both lower and higher bands, the LP-delta clusters reflected mean evoked power decrease for mutual versus unreciprocated cooperation outcomes (Figure 3). No statistically significant clusters were found in the DC versus DD contrast.

**Figure 2.**
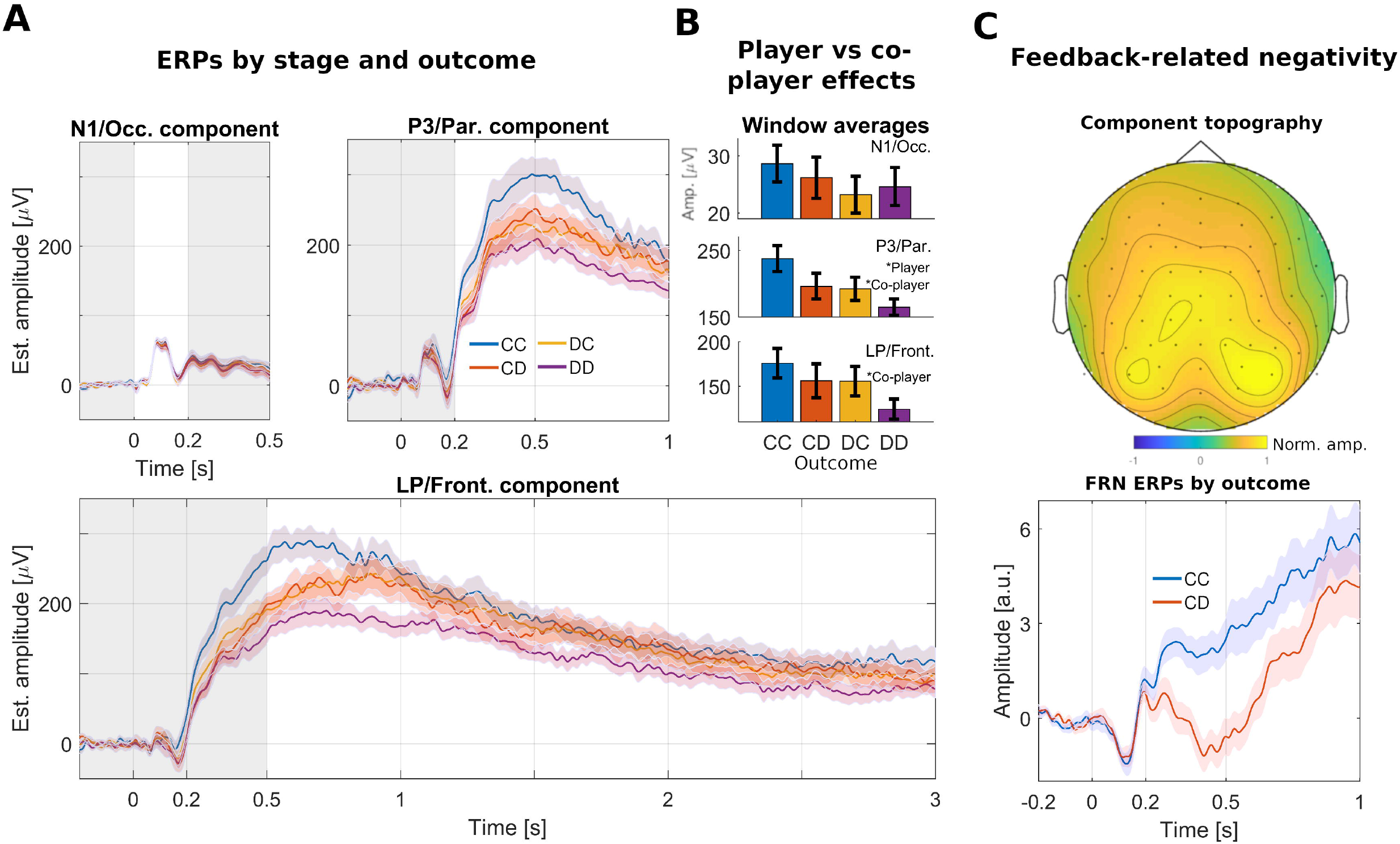
Feedback signals involved in iPD feedback processing. (A) Top left: The visual N1 (0-200 ms; non-shaded region) shows no effects of outcome processing related to players’ choices. Top right: The P3 is the first general feedback signal where effects of player and co-player choices are observed. Bottom: The LP shows an effect of co-player’s choice only. Curve shades indicate ±1 SEM. (B) Summary of window averages and effects found for all general feedback signals and outcomes. (C) The outcome-specific FRN signal was estimated after reproducible scalp activity which differentiates between CC versus CD outcomes across participants. Top: resulting centro-parietal topography of the FRN. Bottom: FRN timeseries at the 0.2 to 0.5 s window shows outcome sensitivity to co-player’s choice given cooperation by the participant.

**Figure 3.**
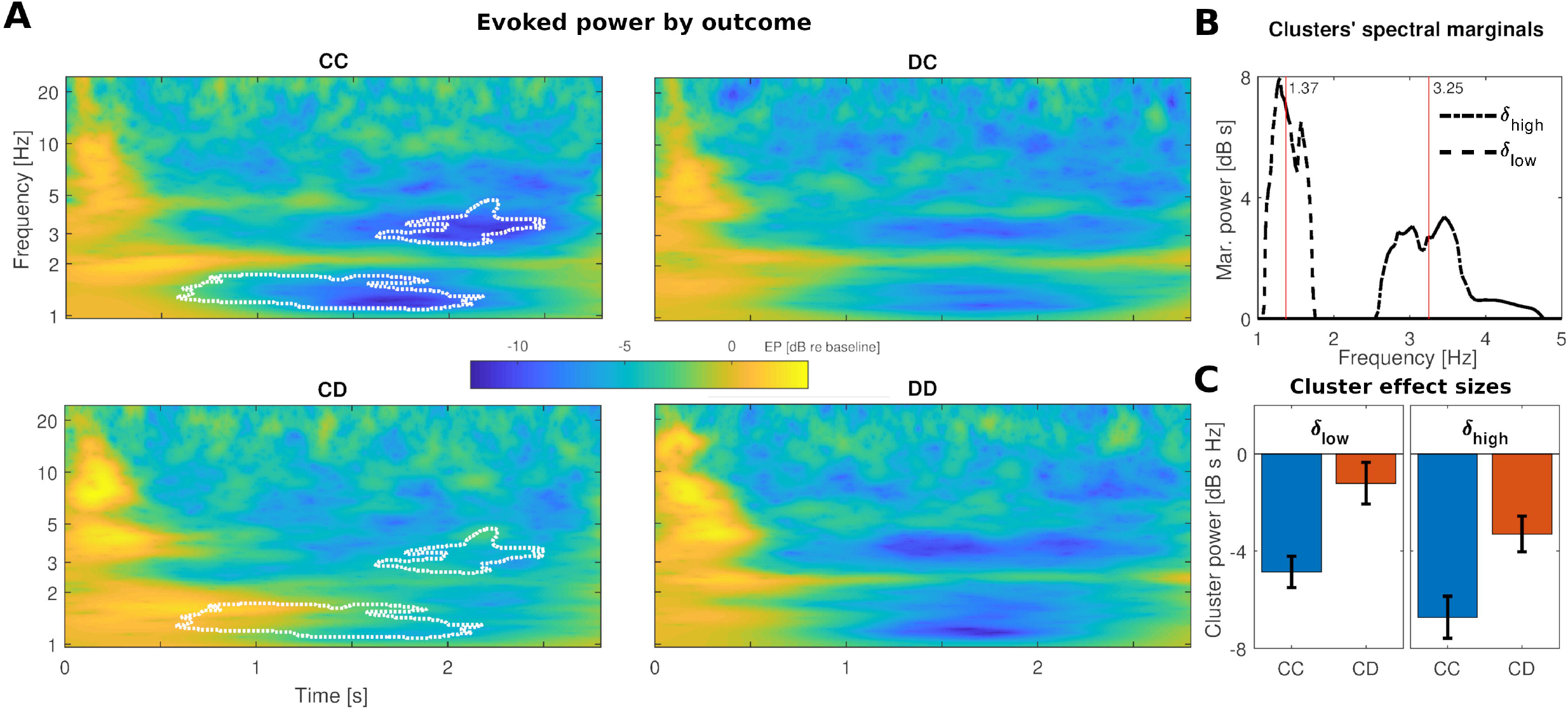
Time-frequency decomposition of the LP feedback signal. (A) Spectrotemporal wavelet correlograms per outcome reveal differential slow-wave (<5 Hz, delta) activity in the 0.5 – 2.5 s interval post feedback onset. Cluster analyses for CC-CD contrasts show significant differences in two time-frequency regions, slow (δ_low_, 0.5-2 s) and fast (δ_high_, 1.5-2.5 approx.) delta-band rhythmic activity (*p* 0.041). Cluster boundaries are superimposed. No clusters were found for DC-DD contrast. (B) Cluster spectral profiles are predominantly contained in the delta band. (C) Effect sizes, defined as participants’ evoked power integrated over relevant cluster region boundaries, show significant effects of cooperation by the co-player, indicating relatively greater suppression for CC outcomes.

### EEG outcome signals indicate future choice and speed of cooperative reciprocation at the iPD

Given their role in outcome processing, FRN, P3 and LP waveforms, plus the LP-delta clusters, were further addressed for the hypothesis that they may be early indicators of participants’ *next choice* to cooperate. For these analyses, next-trial choices were separated by their leading outcome types (e.g., cooperation/defection given mutual cooperation ‘C|CC’ / ‘D|CC’). If a given feedback signal was found to be indicative of next choice, we were then also interested in whether its trial-by-trial amplitude would also relate to the speed to which a given choice was made. In such case, the time to reach a certain choice (e.g. ‘C|CC’) was compared against the amplitude of the relevant feedback signal in the prior trial. For this, repeated measures Pearson correlation coefficients^75^ were computed to test the relationship between reaction time and single-trial signal amplitudes.

#### Mutual cooperation (CC)

Following mutual cooperation, no significant modulations in the feedback FRN, P3 or LP waveforms were found, in relation to next-trial choice. For LP-delta, C|CC trials showed reduced activity compared to D|CC trials (Figure 4A, bottom), within the high-delta region determined earlier. For the spectrotemporal cluster shown by non-parametric testing (half-maximum 3.6-4.3 Hz, 1.87-2.23 s; *p*=0.034), LP-delta was relatively more attenuated in C|CC compared to D|CC (*t*(15)=2.78;*p*=0.014; *d* = 0.70) (Figure 4B). For the low-delta region, no significant modulation by next choice was found. Single-trial LP-delta activity at CC outcomes was further examined in relation to next reaction time, separately for the C|CC and D|CC choices. The data revealed a significant relationship between single-trial LP-delta activity at CC and next trial reaction time to reciprocate (i.e., C|CC), where relative LP-delta decreases were correlated with faster cooperation (*ρ_rm_*(624) 0.084, CI [-0.001, 0.161], *p*=0.035; Figure 4C). No significant correlation was found between single-trial LP-delta and reaction time at D|CC choices.

**Figure 4.**
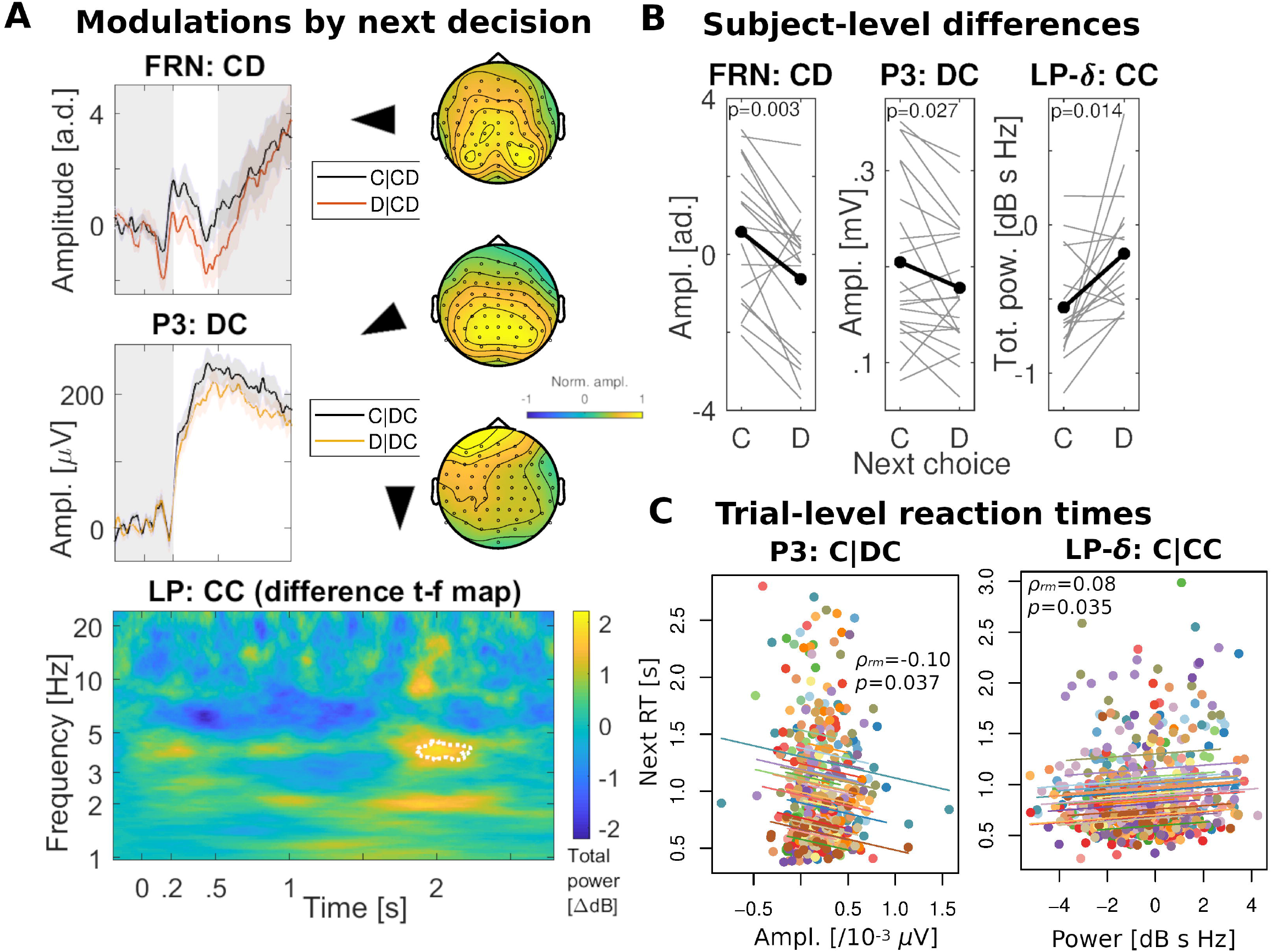
Future round decision-making at the iPD, indicated by scalp activity related to current feedback processing in an outcome-contingent manner. (A) Top: At CD outcomes (sucker’s payoff), centro-parietal FRN relates subsequent round decisions. Non-cooperation was indicated by lower-amplitude FRN signals. Middle: For DC (one-sided non-cooperation), higher P3 amplitudes indicate forthcoming cooperation. Bottom: At CC (mutual cooperation), LP-slow-wave activity also relates next round choice. The difference map in the total power correlogram (D|CC minus C|CC) is shown, with the significant cluster boundary superimposed. Topographies for each component are displayed. (B) FRN, P3 and LP-delta subject-level data (in grey) indicate next round decision-making contingent upon CD, DC and CC outcome types, respectively. Grand average values are shown in black. (C) In the specific decision *to reciprocate co-player’s cooperation* (i.e. C|CC,DC), trial by trial variations in relevant feedback signals (P3 amplitude for DC, LP-delta suppressions for CC) relate how fast individuals will respond at the next trial.

#### Unreciprocated cooperation / sucker’s payoff (CD)

At the CD feedback stage, FRN signals were significantly modulated by the choice to cooperate at the next trial (*t*(17)=3.53; *p*=0.003; Cohen’s *d*=0.83), where D|CD trials had lower FRN amplitudes than C|CD on average (Figure 4A, top). There was limited evidence that the P3 and LP may be significantly modulated by subsequent choice (P3: *t*(17)=2.09; *p*=0.052; *d*=0.49; LP: *t*(17)=1.88; *p*=0.078; *d*=0.44). No LP-delta power cluster differences were significant. At the trial level, FRN activity at CD outcomes was further examined in relation to speed of choice, separately for the C|CC and D|CC cases, but no significant correlations were found.

#### One-sided non-cooperation (DC)

One-sample *t-*tests showed significant mean waveform differences reflecting greater P3 amplitudes at the feedback stage for C|DC than D|DC (*t*(20)=2.39; *p*=0.027; Cohen’s *d*=0.52; Figure 4A,B). Trial P3 activity at DC outcomes was further examined in relation to speed of choice, separately for the C|DC and D|DC cases. Single-trial P3 amplitudes involving C|DC choices correlated negatively with the subsequent reaction time, (*ρ_rm_*(422)=-0.101, CI [-0.196, −0.001], *p*=0.037) (Figure 4C). No significant correlation was found between single-trial P3 and reaction time at D|DC choices.

#### Mutual defection (DD)

No significant mean waveform differences were observed for P3 nor LP signals relating subsequent choice after a DD outcome.

Randomization tests were performed to verify that findings were not a result of systematic bias explainable by sample size imbalances across conditions. Single trial data were randomly shuffled across next decision conditions, per participant, feedback signal, and outcome type (FRN:CD; P3:DC; LP-delta:CC). The observed effects at the FRN (*p*=0.018), P3 (*p*=0.026) and LP-delta (*p*=0.002) regarding modulations by next choice, were found to be significant.

## Discussion

This study examined neural processing at the iPD outcome stage in order to provide a more comprehensive picture of relevant feedback signals, while their ability to indicate subsequent decisionmaking was further investigated. In line with previous studies, FRN and P3 sensitivity to cooperative choice feedback was observed. In addition, inspection of the previously unaddressed late latency period revealed a frontally-distributed component (‘LP’), of which delta-band activity (‘LP-delta’) was also indicative of cooperative choice feedback. Crucially, the feedback signals were, under specific circumstances, also indicative of choice at the next trial: (i) at unreciprocated cooperation (CD), downmodulations of the FRN related to forthcoming non-cooperation; (ii) at one-sided defection (DC), greater P3 amplitude was associated with player’s next cooperation; (iii) at mutual cooperation (CC), LP-delta power reductions similarly related to cooperation at the next round. Single trial P3 activity (at DC rounds) and, separately, LP-delta deactivation (CC) predicted shortening of the time *to cooperate* at the next round. Both scenarios reflect reciprocal reaction, given that the co-player had cooperated. We next discuss the role of each feedback component, and interpret them in relation to relevant social judgments that follow after each associated outcome type. The timing findings may provide a neurally informed account of fast-mode or intuitive cooperative decision-making.

### FRN

As expected, the data showed differential processing between mutual and unreciprocated cooperation for the FRN, with decreases for outcomes of unreciprocated cooperation (CD) relative to mutual cooperation (CC). This is consistent with findings from studies using reward, as well as social tasks, where negative stimuli are associated with incremented deflections of the FRN. This component is considered the earliest index in reward outcome evaluation^27,76,77^, with sources including the anterior cingulate cortex (ACC)^28,69,76^, posterior cingulate cortex (PCC)^29,76,78,79^ and basal ganglia^31^. The observed centroparietal distribution appears consistent with PCC activation (cf.^77,79,80^) a region with frontal connectivity required for cognitive and behavioral control^80,81^.

Feedback-related activity of the PCC during economic games can persist at subsequent decision-making stages, and predict behavior and strategy change therein^69,78,80,82–84^ regarding future action by a co-player^85–87^. Accordingly, in analyses of next decision we found that FRN downmodulations by unreciprocated cooperation (CD) were more prominent if, on the next trial, the participant played noncooperation. FRN modulations are thought to be underlain by dopaminergic firing pauses^77^ and interpreted as a reward prediction error^32,88^ or event surprise or saliency^33,78,89,90^. Findings of equal-payoff CD outcomes leading to alternative choices, may raise the question of whether some CD events are experienced as more of a loss than others, either in social and/or emotional terms.

### P3

The data featured the P3 as a feedback signal that appeared across outcome types in the iPD, but with relative increases for cooperative choices. These results are in line with findings of P3 increments for desired or positive outcomes at social dilemmas such as the chicken^35,38^ and ultimatum games^36,37^, however, they contrast with mixed results for the dictator^91^ and modified versions of the ultimatum^92^ and prisoner’s dilemma^25,26^. In a one-shot analysis of feedback signals at the PD game, cooperative choice by a co-player did not show P3 amplitude changes^25^; in an iterated and stochastic version of the game, P3 increases were shown for co-player non-cooperation outcomes^26^. The relative inconsistency across these findings may suggest that magnitude changes in the P3 are sensitive to the relevant payoff and decision-making structure of a task. General frameworks from the perceptual and value-based decision making literature posit P3 change to reflect the accumulation dynamics of relevant feedback evidence, for communication over decision-making networks^93–100^. The P3 is also linked to evidence integration^101–103^, event categorization^94^, and active working memory dynamics^104–108^. In this line, the variety of effects across studies may be consistent with “temporal filtering” by dedicated attentional and memory resources for outcomes requiring long-term selection for adaptive choice^105,109,110^, e.g. increased likelihood of sustaining benefits. Here, participants were set to maximize points as a goal, under a stochastic tit-for-tat strategy, and the temporal filtering model would predict that (compared with other outcomes) mutual cooperation events across the session deserve careful realization, since they directly increase the prospects of accumulating reward. This interpretation is consistent with the pattern we observed in our findings. In a naturalistic setting where participants may be continually asked on their belief if an opponent will cooperate^26^ (and therefore such assessment acquires especial relevance for performance) one may expect an alternative pattern, where failures to predict cooperation entail maximal P3 change - again consistent with P3 as a feedback signal mediating adaptive choice.

We also found, in next decision analyses, that P3 amplitudes after DC outcomes were more prominent if on the next trial the participant played cooperation. DC outcomes here represent a social context after which decisions require estimation of the co-player’s likelihood to reciprocate after having suffered a loss. In the temporal filtering model perspective, the results may suggest an adaptive dissociation between the processing of certain DC events versus others. All DC scenarios provide with the potential to engage on sustained mutual cooperation and, to that extent, the observed relative differences may indicate dedicated attentional or memory resources where the player chooses to engage cooperatively. The P3 may integrate contributions from social processing systems, including theory of mind, working memory and planning processes, that are subserved across parietal networks^111–113^. A potential interpretation of the findings, in line with the previous results for the FRN at CD outcomes, is that P3 change by next decision reflects differential surprise or outcome saliency computations across DC events. For the FRN, this surprise was referenced to saliency experienced by the player herself; a possibility is that in DC outcomes, saliency experimented from the co-player-perspective may be estimated^91^. Finally, the P3 feedback latencies observed for DC scenarios could raise questions on conditions for access to conscious choice^114,115^.

### LP/LP-delta

We observed a late frontal evoked signal (‘LP’) that was sensitive to co-player choice. In terms of latency, this component appears consistent with the late positive potential (‘LPP’), a feedback signal related to emotional processing. However, on a spatial basis, the LPP typically follows a posterior scalp distribution associated with visual cortical areas^27,54^ (see ^56^ for a more frontal distribution). The LPP often displays a positive bias for negative stimuli or losses in value-based decision making studies^27,54–56^, although in emotion studies evidence of negative bias at late latency windows is mixed (cf.^53,111,117^) with converging evidence suggesting arousal indexing from motivational systems^118,119^. The observed LP here, nevertheless, may show overlap with studies that relate slow wave responses of extended duration in relation to sustained attentive processing^53^, and to the orienting response in a motivationally salient context^120,121^.

Waveform signals with such long-term dynamics as the observed LP may motivate spectrotemporal analyses (e.g.^*57*^) given their potential to be driven by slow-wave components; indeed, the results showed a so far unobserved suppression of LP-delta band activity for mutual versus unreciprocated cooperation. A central role of delta-band modulations has been proposed in terms of homeostatic and autonomic processing of reward and emotion cortical systems^122,123^. Delta decrements are considered to reflect more relaxed (or less anxiogenic) states^124^, positive options or feedback^125,126^, while increases may be elicited by arousal^57,123,127^ or during highly focused states^128^ (but see^129^). Arguably, in the iPD, CC outcomes induce less cognitively-demanding states because they imply a default frame for both players (e.g.^130^). Our findings of delta band suppression during mutual versus unreciprocated cooperation appears consistent with more relaxed states induced by the reciprocal and advantageous outcome.

Moreover, we observed that frontal LP-delta suppressions after mutual cooperation were more pronounced as players later chose to maintain cooperation. As our findings relate to a frontal scalp distribution, the delta-band sources may, in principle, involve areas such as medial and ventral frontal cortex^131–134^ known to modulate delta activity^135^. Such frontal areas signal upcoming decisions following feedback presentations^136^, and are specialized for planning and control (e.g.^137,138^). They additionally involve systems that represent game option values^139–143^ and/or socio-emotional perspective taking^144,145^. Since delta activity changes may be related to self-adjusting mechanisms of homeostasis, it is an open question whether the observed effects in next decision analyses reflect distinct representations of value in homeostatic terms^126^, or anticipated cooperation risk/belief (e.g.^26^). Future studies may aim to clarify whether such deactivations represent value signals or updating processes^146^.

#### Faster cooperative decision-making at the iterated PD

With regards to timing, P3 (after DC outcomes), and LP-delta (after CC) were the only feedback signals indicative of single-trial reaction times, if participants would cooperate at the next round. In behavioral analyses, we found faster responses by around 70 ms after CC than DC outcomes. On the other hand, EEG analyses showed that, at CC, LP-delta feedback signal modulations by subsequent choice took considerably longer to appear (~1.5 s) relative to analog P3 effects at DC. The relative differences potentially suggest distinct timelines in cooperative decision-making, with a comparatively shorter decisional process for the specific action *to cooperate after a mutual cooperation* than after a one-sided non-cooperation. In the SHH^63^ framework, cooperative strategies that are advantageous in routine interactions may become ‘internalized’ towards automatic and/or emotional dispositions, because doing otherwise is relatively slower, effortful and potentially signals indecision. Our results of shorter processing in cooperating after CC compared to DC suggest a greater role of intuitive cooperation at CC conditions. After DC, cooperating may entail deliberation on how trustworthy is the co-player given the unadvantageous outcome.

With regards to timing, P3 (after DC outcomes), and LP-delta (after CC) were the only feedback signals indicative of single-trial response times, provided participants would cooperate at the next round. Neural signals’ magnitude at a given round hence served to predict speed of cooperative reciprocation at the next. In this regard, faster choices were individually associated to pronounced neural dynamics changes, as soon as at the feedback stage of the previous round. The results suggest that automatic or intuitive decision-making may be predicted as soon as during early neural signaling that indexes learning of a co-player’s *cooperative* choice. Moreover, the dynamics at this learning stage may further predict the speed at which such action is reciprocated at the next opportunity, possibly involving attentional and homeostatic signaling. In the case of *reciprocating mutual cooperation*, both behavioral and neural results provide the strongest support for the involvement of fast-mode or intuitive decision-making^60–62,147^, with frontal delta deactivation as its neural signature of autonomic function.

## Conclusion

A comprehensive description of neural signals involved in the iterated prisoner’s dilemma is crucial to identify the neural basis of cooperative strategies. We extend previous ERP literature to include long latency but also slow-wave activity among the feedback processes that index present but also future cooperative choices. Each outcome type may entice alternative predictive strategies according to the socially learned situation at present^146,148^, which was reflected in findings that feedback signals encoding cooperative choices also relate future cooperation, in an outcome-dependent manner. For the specific action of *reciprocating mutual cooperation*, future expedited executions were associated to delta-band activity that has been associated to homeostatic processing in the literature. The results support the involvement of cognitive systems underlying fast decision-making^59,147^, relating them to the decision to cooperate in a social dilemma^60–62^. We suggest a direct involvement of such neural signaling with social heuristics deployment.

## Acknowledgements

This work was supported by *Proyectos I+D* from the Comisión Sectorial de Investigation Científica (UdelaR), and *Fondo Santiago Achúgar Díaz* 2018 (Uruguay) to V.B.G. We thank support from the Agencia Nacional de Investigation e Innovación (Uruguay) to F.C.C., and from *Ayudas a la Atracción de Talento Investigador* and *Proyectos I+D para jóvenes investigadores*, Comunidad de Madrid and Universidad Autónoma de Madrid (Spain) [2017-T2/SOC-5569; SI1-PJI-2019-00011]. We also thank funding support from ANII and Programa de Desarrollo de las Ciencias Básicas to A.C. and V.B.G

## Data availability statement

The data that support the findings of this study are openly available in Open Science Framework at http://doi.org/10.17605/OSF.IO/G8Z7Y

