## Supplementary Data / Figures for "Neural processing of iterated prisoner’s dilemma outcomes indicates next-round choice and speed to reciprocate cooperation"

### Appendix - Supplementary Data

<u>*Emotional and behavioral data analyses*</u>
Emotional Likert scores were normalized from 0 ('not at all' experienced) to 1 ('extremely') and analyzed with a two-way ANOVA with emotion and outcome type as factors. Reaction times were computed as the interval between payoff matrix presentation and player button press, and computed per preceding outcome (Figure 1B). Transition probabilities, (i.e. the probability that the player will cooperate at the next trial given the current outcome) and reaction times were analyzed with a two-way ANOVA with Player and Co-player choices as factors.

#### *Emotional and behavioral results*

iPD outcomes triggered emotions and behaviors as expected (Figure 1A, Supplementary Figure 1C). A 2-way repeated measures ANOVA on emotional ratings identified significant main effects for emotion type ($F(4,116)=68.1$; $p<0.001$; $\eta_p^2=0.701$), outcome type ($F(3,87)=9.8$; $p<0.001$; $\eta_p^2=0.252$), and a significant emotion by outcome interaction ($F(12,348)=29.6$; $p<0.001$; $\eta_p^2=0.505$). Each outcome type was associated with specific emotional reactions (Figure 1B), consistent with previous work[5,15]. CC outcomes were associated with the emotion of happiness; CD with anger, sadness and betrayal; DC with satisfaction and guilt; and DD with intermediate levels of all emotions.

Behaviorally, reaction times and probability of cooperation following each outcome type were analyzed. A 2 x 2 factorial repeated measures ANOVA with Player and Co-player choices as independent variables showed a significant main effect of Player choice on subsequent reaction time ($F(1,28)=5.64$; $p=0.025$; $\eta_p^2=0.168$) (Figure 1B), and no significant effect by Co-player choice ($F(1,28)=1.84$; $p=0.19$; $\eta_p^2=0.06$). The main effect was qualified by a significant interaction between Player and Co-player choices ($F(1,28)=4.44$; $p=0.044$; $\eta_p^2=0.14$). Post-hoc comparisons indicated that reaction times were on average 77 ms faster (CI: 28 to 125 ms; Cohen's $d=0.6$) after CC than DC outcomes ($t(28)=3.23$; $p=0.003$). By contrast, CD versus DD reaction times did not significantly differ ($t(28)=0.06$; $p=0.95$).

In terms of decision-making, transition probabilities, (i.e. the probability that the player will cooperate at a next trial given the type of outcome in the present trial) were analyzed (Supplementary Figure 1C). A 2 x 2 factorial repeated measures ANOVA was run with Player and Co-player choices (i.e. C versus D each) as independent within-subjects variables. A significant main effect of Player choice on subsequent probability to cooperate was found ($F(1,29)=24.7$; $p<0.001$; $\eta_p^2=0.460$), where cooperation by the Player led to higher chances of cooperating in the next round. Similarly, a significant main effect of Co-player choice was found ($F(1,29)=17.4$; $p<0.001$; $\eta_p^2=0.376$), where cooperation led to a higher probability that the participant will cooperate in the next round. The interaction between Player and Co-player choices on transition probability was not significant ($F(1,29)=0.52$; $p=0.48$; $\eta_p^2=0.02$). In terms of reaction times, when all trials were pooled and analyzed for the reaction time to make a decision to cooperate or not, a non-significant trend was found in the differences taken to cooperate (880 ± 173 ms; mean ± SD) and to defect (946 ± 197 ms) ($t(28)=2.000$; $p=0.056$).

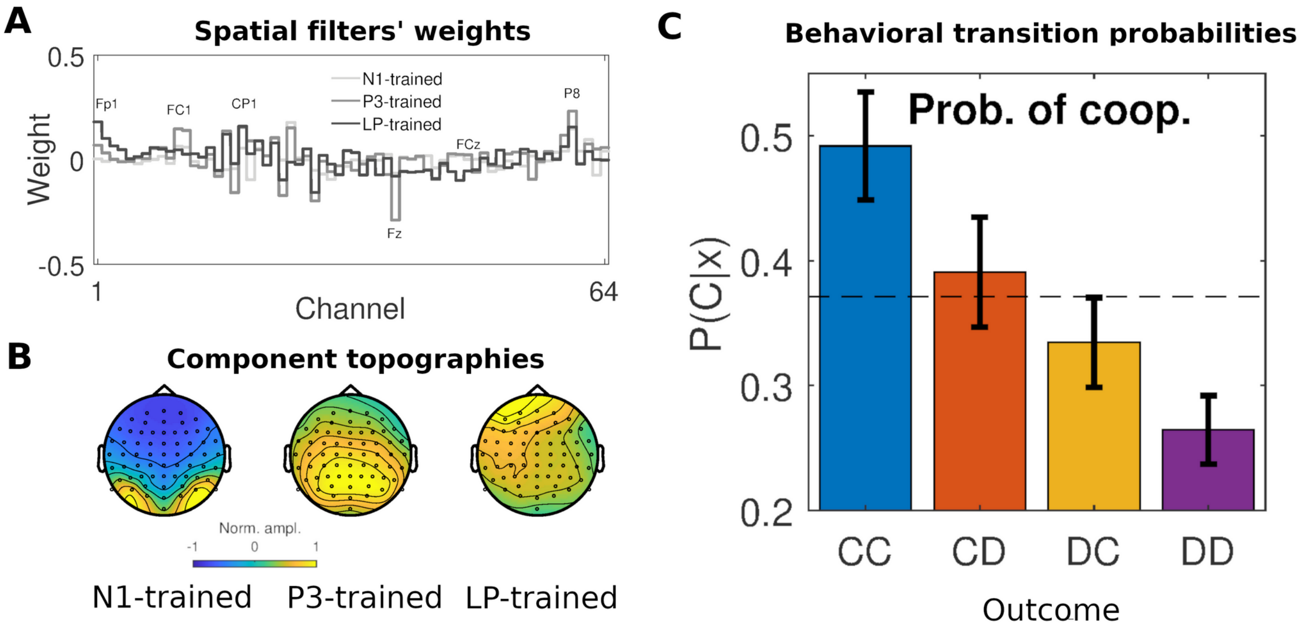


**Supplementary Figure 1.** (A) Spatial filters trained on the basis of N1- (light), P3- (medium) or late potential-epochs (dark). Once estimated, each filter consists of a 64 array of linear coefficients designed to reveal specific components when applied to the whole EEG data. A selection of relevant sensors is shown. (B) Topography of the components associated with each spatial filter, and with reproducible activity across trials/participants. (C) The probability to cooperate after a trial is influenced by the involved decisions in the present trial; dashed line indicates average baseline levels. Error bars indicate ±1 SEM.
